# Functionalized mesoporous silica nanoparticles for innovative boron-neutron capture therapy of resistant cancers

**DOI:** 10.1101/471128

**Authors:** Guillaume Vares, Vincent Jallet, Yoshitaka Matsumoto, Cedric Rentier, Kentaro Takayama, Toshio Sasaki, Yoshio Hayashi, Hiroaki Kumada, Hirotaka Sugawara

**Author notes:** These authors contributed equally (Guillaume Vares, Vincent Jallet). Biotage Japan, Koto, Tokyo 136-0071 (Cedric Rentier). High Energy Accelerator Research Organization (KEK), Tsukuba, Ibaraki 305-0801.

## Abstract

Treatment resistance, relapse and metastasis remain critical issues in some challenging cancers, such as chondrosarcomas. Boron-neutron Capture Therapy (BNCT) is a targeted radiation therapy modality that relies on the ability of boron atoms to capture low energy neutrons, yielding high linear energy transfer alpha particles. We have developed an innovative boron-delivery system for BNCT, composed of multifunctional fluorescent mesoporous silica nanoparticles (B-MSNs), grafted with an activatable cell penetrating peptide (ACPP) for improved penetration in tumors and with Gadolinium for magnetic resonance imaging (MRI) *in vivo*. Chondrosarcoma cells were exposed *in vitro* to an epithermal neutron beam after B-MSNs administration. BNCT beam exposure successfully induced DNA damage and cell death, including in radio-resistant ALDH+ cancer stem cells (CSCs), suggesting that BNCT using this system might be a suitable treatment modality for chondrosarcoma or other hard-to-treat cancers.

## Background

Remarkable progress has been made in the understanding and treatment of human cancer, resulting in a greatly improved survival rate for many patients. However, such achievements remain incomplete or out of reach for some hard-to-treat cancers, such as pancreatic cancer, glioblastoma or bone tumors. Chondrosarcomas are cartilaginous tumors which represent the second most common primary bone tumor in adults [1]. Chondrosarcomas are notoriously resistant to conventional radiation therapy and to chemotherapy, and complete surgical resection remains to this day the primary treatment. A number of patients experience relapse, metastasis or present unresectable disease with poor clinical outcome [2].

Tumors are heterogeneous and are comprised of cells with various morphological and molecular features, including a subset of tumor-initiating dedifferentiated cells with self-renewing abilities. These cancer stem cells (CSCs) are capable of reconstituting tumor heterogeneity, and a large amount of evidence strongly suggests that they may contribute to treatment resistance, relapse and metastasis [3]. CSCs have been identified in a number of tumors, including chondrosarcomas [4]. Several features of CSCs have been reported to explain their intrinsic radioresistance: lower levels of basal and radiation-induced reactive oxygen species (ROS), improved DNA damage repair activation and apoptosis inhibition or relative quiescent state [5]. New approaches are thus highly expected to address treatment-resistant tumors, which may include targeting CSCs.

In addition to conventional X-ray therapy, new high linear linear energy transfer (LET) radiation therapy modalities have emerged which might finally contribute overcoming treatment resistance, such as boron neutron capture therapy (BNCT) or carbon-ion particle therapy [6]. High LET therapies, alone or combined with other targeted treatments (like the PARP inhibitor talazoparib or the chemotherapeutic drug cisplatin), may be able to overcome CSC-related resistance [6–11]. BNCT is an innovative experimental treatment modality that relies on the ability of ^10^B to capture thermal neutrons, resulting in the release of high-linear energy transfer (LET) α (^4^He) particles and lithium (^7^Li) nuclei, with a path length shorter than 10 *μ*m [12]. Therefore, it is crucial to maximize the concentration of boron-enriched compounds in tumor tissues while minimizing levels in surrounding normal tissues. Furthermore, intracellular boron delivery should be achieved, due to the short path length of α (^4^He) particles. Because BNCT releases high-LET radiation, it should provide improved relative biological effectiveness (RBE) and a lower oxygen enhancement ratio (OER), compared to conventional X-ray therapy. BNCT clinical trials have been performed on patients suffering from head and neck, brain, lung and liver cancers [12], with some encouraging results in terms of overall survival, recurrence and metastasis. For those reasons, BNCT might also be an effective strategy for the treatment of radioresistant tumors, such as clear cell sarcoma (CSS) [13], osteosarcoma [14] or chondrosarcoma.

Even though the first BNCT trials have been performed more than half a century ago, BNCT has not yet become an established treatment modality, due to two main limiting factors [15]. First, only two boron-delivery drugs are routinely used in BNCT clinical trial studies: sodium mercaptoundecahydrododecaborate (Na_2_ ^10^B_12_H_11_SH; Na_2_ ^10^BSH) and L-*p*-boronophenylalanine (L-^10^BPA). Reported tumor-to-normal tissue (T/N) ratio for BSH does not always reach 1. New advances are necessary to improve T/N ratio with low toxicity. Second, the sole neutron sources traditionally available for BNCT were nuclear reactors. Recent advances in nanotechnologies for drug delivery and the development of new accelerator-based neutron sources promise to overcome those limitations. Here, we report a new theranostic multi-functional boron-delivery system based on mesoporous silica nanoparticles (B-MSNs). We tested this system using an accelerator-based neutron beam for BNCT.

Nanoparticles (NPs) have recently emerged as a promising therapeutic tool for a variety of medical applications. NPs allow the encapsulation of therapeutic compounds with higher protection against metabolic degradation and the ability to control and target drug release preferentially in tumor tissues [16]. The accumulation of NPs in tumor tissues has been attributed to the poor alignment of neovascularization and lymphatic drainage in those areas, so called enhanced permeability (EPR) effect [17]. In particular, much attention has been devoted to the design of MSNs, which present a number of advantageous features: tunable size (usually 50 to 200 nm diameter), easy surface functionalization, large mesopore volume for efficient drug loading, *in vitro* and *in vivo* tolerance [18,19]. They might therefore serve as ideal boron-delivery agents for BNCT.

## Methods

### Synthesis of mesoporous silica nanoparticles

Synthesis steps are represented in Scheme S1.

#### Synthesis of fluorescent MCM-41 (**1**)

FITC (5.5 mg) was dissolved in 3 mL of ethanol under argon, then APTES (12 μL) was added and the reaction was stirred for 2 hours at room temperature. In a separate flask, CTAB (0.5 g) was dissolved in a mixture of MilliQ water (240 mL) and NaOH 2M (1.75 mL), and the mixture was stirred at 80°C. After 2 hours, the FITC-APTES solution was slowly added to the CTAB solution and the reaction was carried out at 80°C. After 2 hours, the reactional mixture was allowed to cool down to room temperature and the precipitate was filtrated with a fritted glass (porosity 3) and thoroughly washed with methanol. The powder was crushed and dried overnight under high vacuum.

#### Grafting of APTES on nanoparticle surface (**2**)

Nanoparticles obtained during step 1 - NP(**1**) - were suspended in methanol (50 mL) for 5 minutes, then APTES (3 mL) was added and the mixture was stirred overnight at room temperature. The suspension was centrifuged and washed three times with methanol, then allowed to dry for 2 hours under high vacuum.

#### Surfactant removal (**3**)

NP(**2**) were dispersed in a mixture of HCl (5 mL) and methanol (90 mL) and stirred at reflux for 24 hours. The suspension was centrifuged and washed with methanol, then allowed to dry for 2 hours under high vacuum.

#### Ammonium neutralization and aggregates elimination (**4**)

NP(**3**) were dispersed in DMSO (50 mL) then piperidine (12.5 mL) was added to neutralize ammonium. The mixture was bath sonicated for 40 minutes at room temperature, then centrifuged at 500 g for 10 minutes to eliminate aggregates. Nanoparticles were centrifuged and washed twice with DMSO and twice with methanol. The nanoparticles were dried under high vacuum for 2 hours.

#### Grafting of MeO-PEG-COO-NHS (5 kDa) and Mal-PEG-COO-NHS (10 kDa) (5)

NP(**4**) were dispersed in DMSO (50 mL), then Mal-PEG-NHS 10 kDa (5 mg) was added to the nanoparticles and stirred for 30 minutes. Next, MeO-PEG-NHS 5 kDa (95 mg) was added to the nanoparticle suspension. The mixture was stirred over night at room temperature, then centrifuged and washed three times with methanol. The nanoparticles were dried overnight under high vacuum.

#### Grafting of APTES inside pores (**6**)

NP(**5**) were dispersed in methanol (50 mL) for 15 minutes in sonication bath, then APTES (3 mL) was slowly added. The mixture was stirred for 24 hours at room temperature then the suspension was centrifuged and the nanoparticles are washed three times with methanol. The nanoparticles were dried overnight under high vacuum.

#### Grafting of HOOC-CH2-Br inside the pores (**7**)

NP(**6**) were dispersed in DMSO for 15 minutes in sonication bath, then N,N’-Dicyclohexylcarbodiimide (DCC) and N-hydroxysuccinimide (NHS) were added to the nanoparticle suspension. Finally, HOOC-CH2-Br was added and the mixture was left reacting over night at room temperature. The nanoparticles were centrifuged and washed three times with methanol. The nanoparticles were dried under high vacuum for 2 hours.

#### Incorporation of BSH inside pores (**8**)

NP(**7**) were dispersed in a mixture of ACN/H2O (50 mL in proportion 4/1) under argon, then ^10^B-enriched Sodium mercaptoundecahydro-dodecaborate (Na_2_[^10^B_12_H_11_SH]) (Stella Chemifa, Osaka, Japan) (250 mg) and TCEP were added (300 mg). The reaction was stirred under argon overnight at room temperature. The suspension was centrifuged, and the nanoparticles are washed three times with methanol. The nanoparticles were dried under high vacuum for 2 hours. m = 290 mg

#### Grafting of Activatable cell penetrating peptide (**9**)

NP(8) were dispersed in PBS (15 mL) adjusted to pH 7.0 with HCl. In a separate flask, Activatable cell penetrating peptide (ACPP) (1 mg) and tris(2-carboxyethyl)phosphine (TCEP) (6.6 mg, 100 equivalents) were dissolved in PBS pH 7.0. The pH was adjusted from 3.5-4.0 to 7.0 with 1M NaOH. Argon was bubbled in the nanoparticle suspension maintained in a sonication bath for one hour, and the peptide solution was bubbled with argon for 1 h. Then, the peptide solution was slowly added to the nanoparticle suspension and stirred for one additional hour at room temperature under argon. The mixture was centrifuged and washed twice successively with PBS, water and DMSO. Then, the nanoparticles were suspended in DMSO (30 mL) and bath sonicated for 1 hour at room temperature, in order to disaggregate the nanoparticles. Aggregates were removed by centrifugating the nanoparticles at 500 g for 10 min, then the nanoparticles were centrifugated at 30,000 g for 30 min and washed 3 times with methanol. The nanoparticles were dried overnight under vacuum.

##### DLS and zeta-potential

Nanoparticles suspensions in milliQ water (100 *μ*g/mL) were analyzed on a Zetasizer Nano ZS instrument (Malvern Instruments, Malvern, UK). Hydrodynamic radius and zeta-potential were analyzed by Dynamic Light Scattering (DLS) and Laser Doppler Velocimetry (LDV), respectively.

##### Synthesis of Activatable Cell Penetrating Peptide (ACPP)

Reagents and solvents, which were used as received, were purchased from Wako Pure Chemical Industries (Osaka, Japan), Sigma-Aldrich (St. Louis, MO), Watanabe Chemical Industries (Hiroshima, Japan), and Tokyo Chemical Industries (Tokyo, Japan). Activatable Cell Penetrating Peptide (ACPP) Ac-E_8_PLGLAGR_8_N-Acp-C-NH_2_ was synthesized using the Fmoc (9-fluorenylmethyloxycarbonyl) / *t*Bu (*tert*-Butyl) based solid-phase method. Standard protected Fmoc-amino acids (0.141 mmol) or Fmoc-ε-Acp-OH (N_ε_-(9-fluorenylmethyloxycarbonyl)-6-aminohexanoic acid) were sequentially coupled to a Fmoc-NH-SAL Resin (100 mg, 0.047 mmol) using the DIPCI (0.141 mmol)-HOBt (0.141 mmol) method. Coupling steps were performed for 2 h in DMF (1.0 mL) after removal of each Fmoc group with 20% piperidine-DMF (1.5 mL, 5+15 min) to obtain resin-bound protected peptide. N-terminal acylation was carried out by reacting the peptide resin at r.t. with acetic anhydride (Ac_2_O) / *N,N*-diisopropylethylamine (DIPEA) (50 eq + 50 eq) in DMF for 1 hour. Cleavage from the resins and removal of side chain protecting groups were achieved by treatment with TFA-*m*-cresol-thioanisole-EDT (4.0 mL, 40:1:1:1 v:v:v:v) for 3 h at room temperature, followed by preparative reversed phase (RP)-HPLC purification in a 0.1% aqueous TFA-CH_3_CN system to obtain the desired ACPP as TFA salts. The purity of synthesized ACPP was > 95% in RP-HPLC analysis using a SunFire C18 reverse-phase column (4.6 x 150 mm, 5μm) (Waters, Milford, MA, USA) with a binary solvent system: a linear gradient of CH_3_CN (10-30%, 40 min) in 0.1% aqueous TFA at a flow rate of 1.0 mL/min, detected at UV 230 nm. Yields of ACPP obtained as a white powder were calculated as TFA salts. High-resolution mass spectrometry (HR-MS) (TOF MS ES+) was recorded on a Micromass LCT (Waters). Results were: Yield: 51.3% (42 mg); HRMS m/z [M+H]^+^ found: 3179.6292 (calculated for C_127_H_219_N_51_O_43_S: 3179.6317); HPLC purity: 96% (t_R_ = 17.79 min) (Figure S2).

##### Inductively coupled plasma mass spectrometry (ICP-MS)

B-MSNs were dissolved at a concentration of 1 mg/mL in 1M Potassium hydroxide (KOH), then kept in an ultrasound bath for 24 hours. Then MilliQ H_2_O and 5% of Nitric acid (HNO_3_) 67% were added. Determination of boron content in the samples was performed using an Element 2 ICP-MS system (Thermo Scientific, Waltham, MA, USA). No glass container or glass equipment were used for ICP-MS experiments, to avoid sample contamination.

##### Transmission electron microscopy (TEM)

Nanoparticles were suspended at a concentration of 0.2-1.0 mg/mL in water, and 20 *μ*L were deposited on a copper grid covered by carbon film. After 30 seconds, the drop was removed with paper tissue and the grid was air dried for 1 hour. Observation of nanoparticles was performed under a JEM1230R electron microscope (JEOL) at the energy of 100 keV. Nanoparticles element composition was analyzed on a JEM-ARM200F electron microscope (JEOL) using energy-dispersive x-ray spectroscopy (EDS) and electron energy-loss spectroscopy (EELS). For *in vitro* visualization of nanoparticles cell penetration, CH-2879 chondrosarcoma cells were treated with nanoparticles, then were prepared according to standard protocols as recommend by the electron microscope manufacturer (JEOL). Briefly, cells were fixed with 2.5% glutaraldehyde in PBS, then postfixation was performed using 1% osmium tetroxide (OsO_4_) in PBS. Fixed cells were dehydrated using increasing concentrations of ethanol (from 50% to 100%). Substitution was performed using various mixes of propylene oxide (PO) and epoxy resin. Cells were then embedded in TAAB EPON 812 epoxy resin, placed in a silicon embedding plate and resin was polymerized at 60°C for one day. 70 nm sections were placed on a TEM grid and observed on a JEM1230R electron microscope (JEOL).

##### Mouse experiments

4-week old BALB/c nu/nu male mice (Japan SLC, Hamamatsu, Japan) were maintained on a 12-hr light/12-hr dark cycle in a temperature-controlled (22°C) barrier facility with free access to water and a normal diet (CLEA Japan, Tokyo, Japan). Mice were allowed to acclimatize for 5 days before the experiment. 10^6^ CH-2879 chondrosarcoma cells mixed 1:1 with Matrigel Growth Factor Reduced (Corning, Corning, NY, USA) were injected subcutaneously into mouse flank. Tumor volumes were measured using calipers. Mice were used for MRI experiments when tumor volume reached at least 500 mm^3^. Mouse experiment protocols were approved by the Animal Care and Use Committee at Okinawa Institute of Science and Technology Graduate University.

##### Magnetic resonance imaging (MRI)

Gadolinium-nanoparticles diluted in sterile PBS were injected into the tail vein of tumor-bearing mice. Mice were scanned at regular intervals under isoflurane anesthesia using an 11.7-T Bruker MRI (Bruker Biospec GmbH, Ettlingen, Germany). T1 maps were carried out using rapid acquisition with relaxation enhancement (RARE) with TE= 6 ms, variable TR= 200, 531, 958, 1557, 2568, 7500 ms, flip angle (FA)= 90-degree, number of averages (NA)= 1. T2 maps were carried out using multiple-spin echo with 60 TEs from 6 to 179 ms, TR= 2200 ms, FA= 90-degree, NA= 1. Imaging planes were coronal slices with FOV= 38 × 28 mm, matrix size= 256 × 96, and slice thickness= 2 mm for both T1 and T2 maps.

##### Cell culture and sorting of cancer stem cells

CH-2879 chondrosarcoma cells [20] and T/C 28AT immortalized chondrocytes [21] were grown in RPMI1640 and DMEM media, respectively (Nacalai, Kyoto, Japan) supplemented with 10% fetal bovine serum (FBS) (Cosmo Bio, Tokyo, Japan) and antibiotic-antimycotic solution (Penicillin, Streptomycin, Amphotericin B. Gibco ThermoFisher, Carlsbad, CA, USA). Cultures were grown in 5% CO_2_ at 95% humidity. As previously [22], ALDH activity in the cells was measured by flow cytometry using the ALDEFLUOR kit (Stemcell technologies, Vancouver, BC, Canada). Cells with low and high levels of ALDH enzymatic activity (respectively ALDH^-^ and ALDH^+^ cells) were sorted using a FACSAria cell sorter (BD Biosciences, Franklin Lakes, NJ, USA). As a negative control, cells were treated with diethylaminobenzaldehyde (DEAB), a specific ALDH inhibitor.

##### Western blotting

CH-8279 cells were lysed with radioimmunoprecipitation assay (RIPA) buffer (Santa Cruz Biotechnology, Dallas, TX, USA). Protein concentrations were determined using the Protein Assay CBB solution (Nacalai) using bovine serum albumin (BSA) as a standard. 20 μg protein were loaded on Any kD Mini-PROTEAN TGX precast gel (Bio-Rad, Hercules, CA, USA). After separation, the proteins were transferred onto a PVDF Membrane (Bio-Rad). Membranes were probed with anti-MMP2 (Cell Signaling Technology, Danvers, MA, USA) and anti-a-tubulin (MBL, Woburn, MA, USA) antibodies. Staining was detected using an HRP-conjugated anti-rabbit secondary antibody and Chemi-Lumi One L assay kit (Nacalai) on a ImageQuant LAS 4000 imager (GE Healthcare, Chicago, IL, USA).

##### Evaluation of nanoparticles cytotoxicity

*In vitro* viability was measured by trypan blue exclusion, using a TC10 automated cell counter (Bio-Rad). To evaluate *in vivo* toxicity, nanoparticles were injected in the tail vein of healthy mice and body weight was measured for several weeks.

##### Radiation exposure

X-ray experiments were performed using an M-150WE X-ray generator (Softex, Tokyo, Japan) at 140 kVp, 8 mA, 80V. Irradiation dose-rate was 1.3 Gy/min. BNCT biological irradiation experiments were performed at the accelerator-based BNCT facility of the Ibaraki Neutron Medical Research Center (Tokai, Japan) [23]. Neutron beam characteristics for each experimental run are summarized in Table S1. 100 *μ*L/mL nanoparticles were administrated 3 hours before irradiation. Shortly before exposure to neutron beam, plated cells were trypsinized and transferred to 0.5 mL microtubes. After irradiation, cells were either processed for immediate analysis or replated for next day analysis, depending on the endpoint.

##### Flow cytometry analysis of apoptosis and DNA damage

Collected samples were analyzed on a Muse flow cytometer (Millipore Sigma, Burlington, MA, USA), using the Muse Multicaspase Assay Kit and the Muse Multicolor DNA damage Kit, according to manufacturer’s instructions.

##### Clonogenic assay

After irradiation, CH-2879 cells and CSCs were seeded in 6-well plates at defined densities, incubated for 10–14 days then stained as previously described[7]. Colonies with >50 cells were scored and surviving fractions were determined after correcting for the plating efficiency.

##### Statistical analysis

Clonogenic survival curve data were fitted to the linear-quadratic model (for X-ray irradiations) or linear model (for BNCT), using the CS-Cal software (www.oncoexpress.de). Statistical significance of the difference between dose-response curves was performed using one-way Analysis of Variance (one-way ANOVA) with Bonferroni correction for pairwise group comparisons, with SigmaPlot software (systatsoftware.com/products/sigmaplot/). Other significant differences were assessed using Student’s *t*-test (p<0.05).

## Results

We have developed multifunctional MSNs, which can serve as boron-delivery carrier and can be monitored for BNCT dosimetry (Figure 1, Scheme S1). The mean diameter of inorganic core, as determined by transmission electronic microscopy (TEM), was around 100 nm (Figure 1b), within the range generally accepted for achieving optimal therapeutic effect [24]: nanoparticles smaller than 10 nm in diameter encounter renal clearance, while bigger nanoparticles may not be able to penetrate and diffuse into the tumor. In order to improve biocompatibility and blood circulation time [25], a layer of polyethylene glycol (PEG, 5kDa 95w%, 10 kDa 5 w%) was grafted onto the surface of nanoparticles by peptide bond (Figure 2a). It is usually considered that grafting of PEG with molecular weight of at least 2 kDa is necessary to avoid clearing by the mononuclear phagocyte system [26]. PEG also allowed steric stabilization of the nanoparticles, which did not form significant aggregates in culture media [27]. After PEG grafting, the hydrodynamic radius of B-MSNs, measured by dynamic light scattering (DLS), was around 200 nm (Figure 2b).

**Figure 1.**
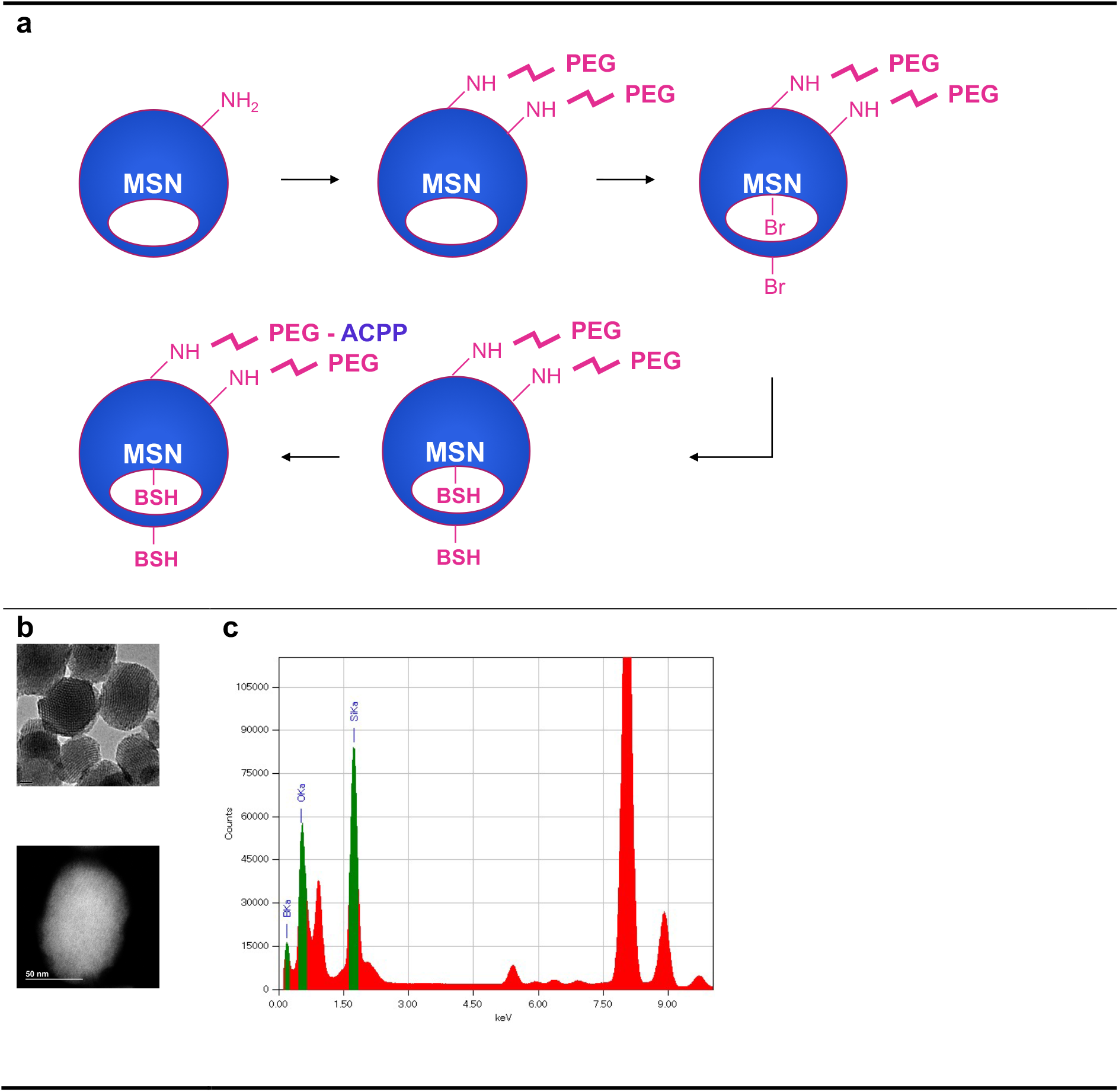
Synthesis of Boron-delivery mesoporous silica nanoparticles (B-MSNs). **(a)** Mesoporous silica MCM-41 fluorescent nanoparticles were PEGylated, then BSH (B10-enriched) was incorporated. Finally, an activatable cell penetrating peptide (ACPP) was grafted. **(b)** Visualization of B-MSNs by transmission electronic microscopy (TEM). **(c)** Energy-dispersive X-ray spectroscopy. Peaks for Boron (BKa), Oxygen (OKa) and Silicon (SiKA) are identified in green.

**Figure 2.**
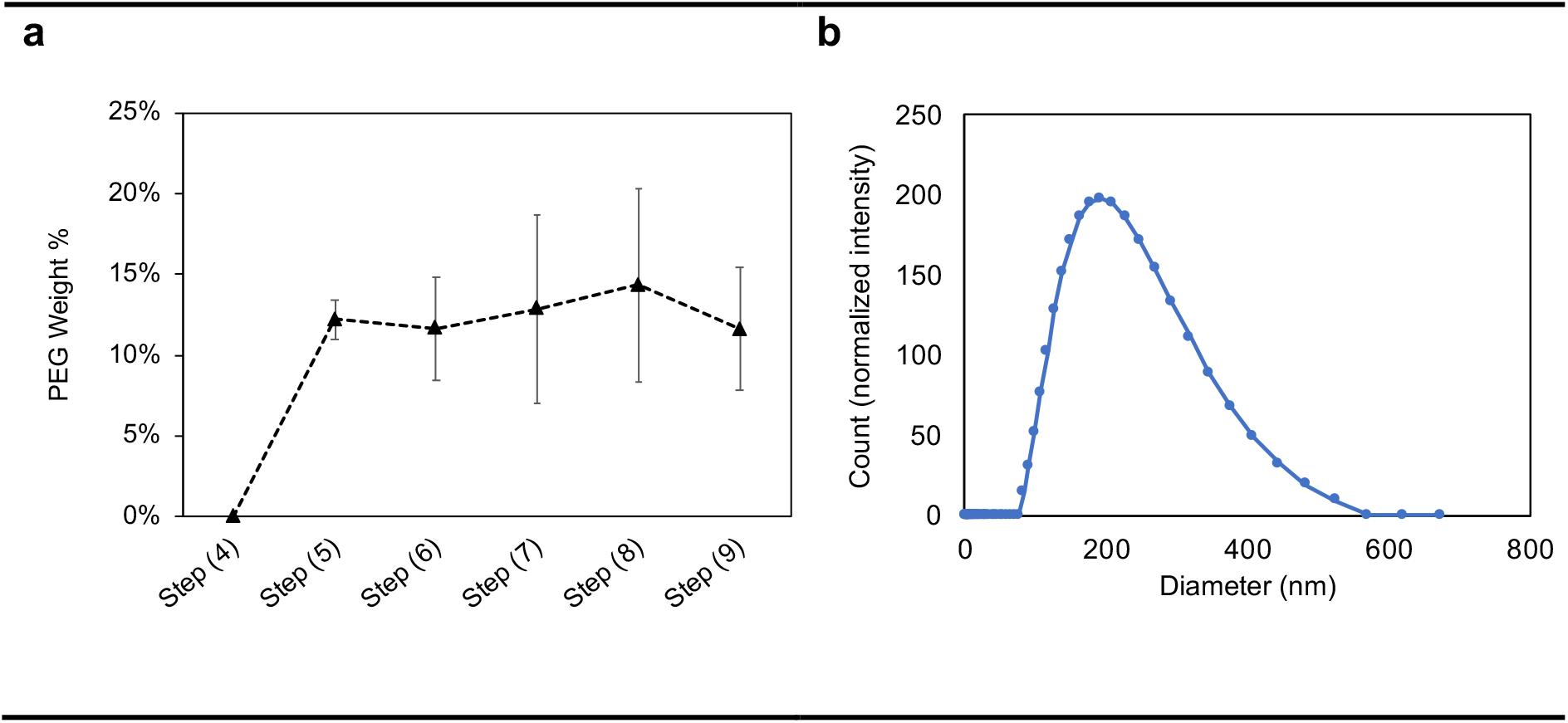
Characterization of Boron-delivery mesoporous silica nanoparticles (B-MSNs). **(a)** NMR quantification of PEG grafted on MSNs at various synthesis steps. **(b)** Measurement of the hydrodynamic radius of B-MSNs by dynamic light scattering (DLS).

The nanoparticles didn’t show any significant toxicity *in vitro* (Figure S3) or *in vivo* (Figure S4) and exhibited good stability and dispersion properties, as confirmed by analysis of zeta potential values (consistently lower than −25 mV). MSNs are considered to be biocompatible, as toxicity can be observed only for high particle concentrations [28].

Sufficient amounts of ^10^B (about 20 μg/g weight or about 10^9^ atoms/cell) need to be delivered to tumor cells for the success of BNCT [29]. Furthermore, because the track of a particles generated by boron-neutron capture is 10 μm at most, it is necessary that a sufficient proportion of ^10^B penetrate inside cells for optimal efficiency. Large amounts of boron might be loaded into nanoparticle mesopores as o-carborane [30], however there is a risk of carborane leakage and unpredictable boron distribution. Here, we propose to attach ^10^B-enriched BSH inside mesopores, using an aminosilane coupling agent. Inductively coupled plasma mass spectrometry (ICP-MS) measurements confirmed the successful accumulation of ^10^B on B-MSNs, which contain 1.27% mass fraction of boron (95% ^10^B), representing around 5 x 10^17^ atoms of ^10^B per mg nanoparticles (Table 1). Subsequent steps of nanoparticle synthesis did not lead to release of BSH. This suggests that if *in vivo* B-MSN delivery to tumors could be optimized, the amount of boron reaching tumor cells might be sufficient for BNCT treatment.

**Table 1.**
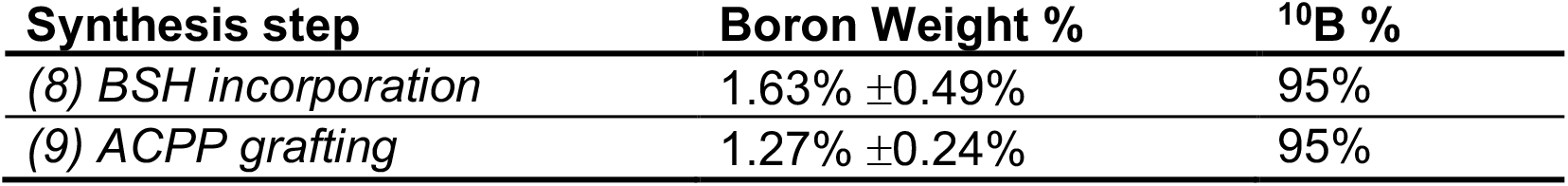
Quantification of Boron content by ICP-MS. Measurements were performed after BSH incorporation and after grafting of ACPP.

| **Synthesis step** | **Boron Weight %** | **<sup>10</sup>B %** |
| --- | --- | --- |
| *(8) BSH incorporation* | 1.63% ±0.49% | 95% |
| *(9) ACPG grafting* | 1.27% ±0.24% | 95% |

In order to efficiently enter cells by endocytosis, nanoparticles are commonly surface-modified with cell penetrating peptides (CPPs). Surface functionalization of the nanoparticles with an activatable cell penetrating peptide (ACPP) allows for efficient tumor targeting. Our ACPP consists of three regions (Figure 3a): a polyanionic autoinhibitory domain (octaglutamic acid E_8_), a PLGLAG linker region (sensitive to proteases) and a cell-penetrating polycationic domain (octaarginine R_8_) [31,32]. In addition, an Acp (aminohexanoic acid) moiety is grafted on the C-terminal portion of the peptide to serve as a spacer between the polycationic domain and the Cys residue for a better efficiency in the thiol-maleimide coupling strategy used. In the intact ACPP, the polyanionic peptide domain prevents uptake of the polycationic domain. Matrix metalloproteinases (MMP) 2 and 9 (generally overexpressed in tumors [33]) cleave the PLGLAG linker, releasing the cell-penetrating R_8_ portion grafted on the nanoparticle. Indeed, chondrosarcoma cells expressed higher levels of MMP-2 than normal chondrocytes (Figure 3b), leading to enhanced relative cellular uptake, compared to nanoparticles grafted only with polyethylene glycol (PEG) (Figures 3c and 3d).

**Figure 3.**
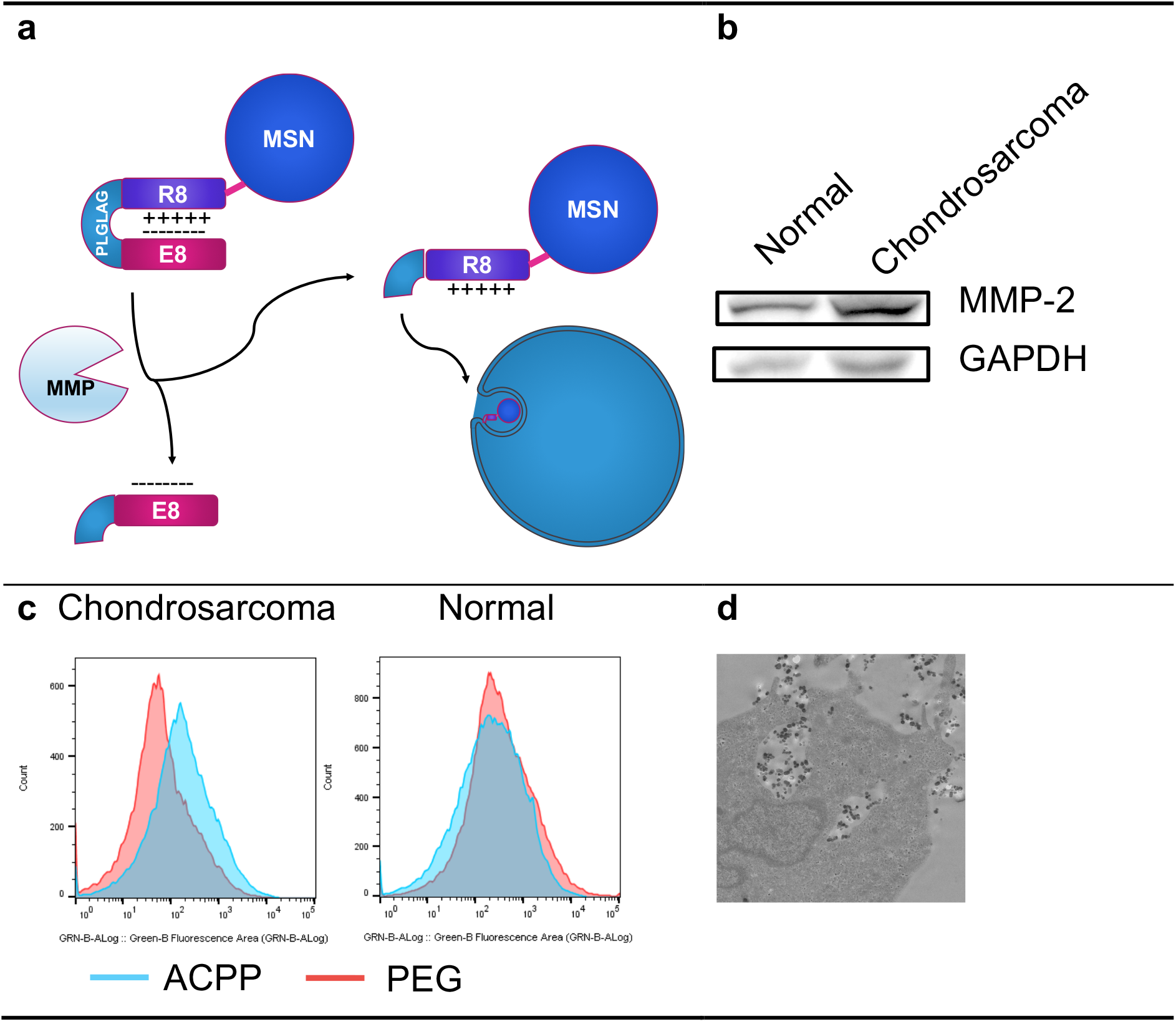
Cellular uptake of functionalized boron-delivery nanoparticles (B-MSNs). **(a)** Matrix metalloproteinase (MMP)-mediated cleavage of the linker PLGLAG region releases the polycationic octaarginine (R8) region of the activatable cell penetrating peptide (ACPP). **(b)** Chondrosarcoma cells express more MMP-2 protein than normal chondrocytes. **(c)** B-MSN cellular uptake is improved in chondrosarcoma cells, as measured by flow cytometry three hours after MSN administration. **(d)** B-MSNs functionalized with ACPP penetrate cells by endocytosis, as observed by transmission electronic microscopy (TEM).

Unlike other radiation therapy modalities (X-rays or charged particle beams), calibration and dosimetry for BNCT relies on many parameters, including neutron beam properties and boron uptake in tumors. In this context, it is crucial to properly monitor the biodistribution of boron-delivery compounds, if possible in a non-invasive way. Although our B-MSNs include Fluorescein isothiocyanate (FITC), allowing fluorescent tracking for *in vitro* and small animal studies, the depth limitation of optical imaging methods seriously hampers their clinical utility. We have therefore also developed MSNs grafted with Gadolinium for *in vivo* visualization using magnetic resonance imaging (MRI) [34,35]. These nanoparticles were injected into the tail vein of nude mice bearing xenograft chondrosarcoma tumors. Longitudinal (*T_1_*) and transverse (*T_2_*) relaxation times were measured for 24h in the tumor (Figures 4, S6). Due to its paramagnetic properties, gadolinium shortens *T_1_* and *T_2_* when it accumulates. While *T_1_* values did not change significantly after injection, we observed a clear decrease in *T_2_* values, reflecting nanoparticles tumor uptake. Relative lack of *T_1_*-weighted contrast is expected at high magnetic fields (11.7 T), as reported previously [36]. Accordingly, increased loading of gadolinium on nanoparticles led to changes in *T_2_*, but not *T_1_* values (Figure S5). Better overall *T_1_*-weighted contrast may be expected at lower fields in MRI systems for routine clinical use.

**Figure 4.**
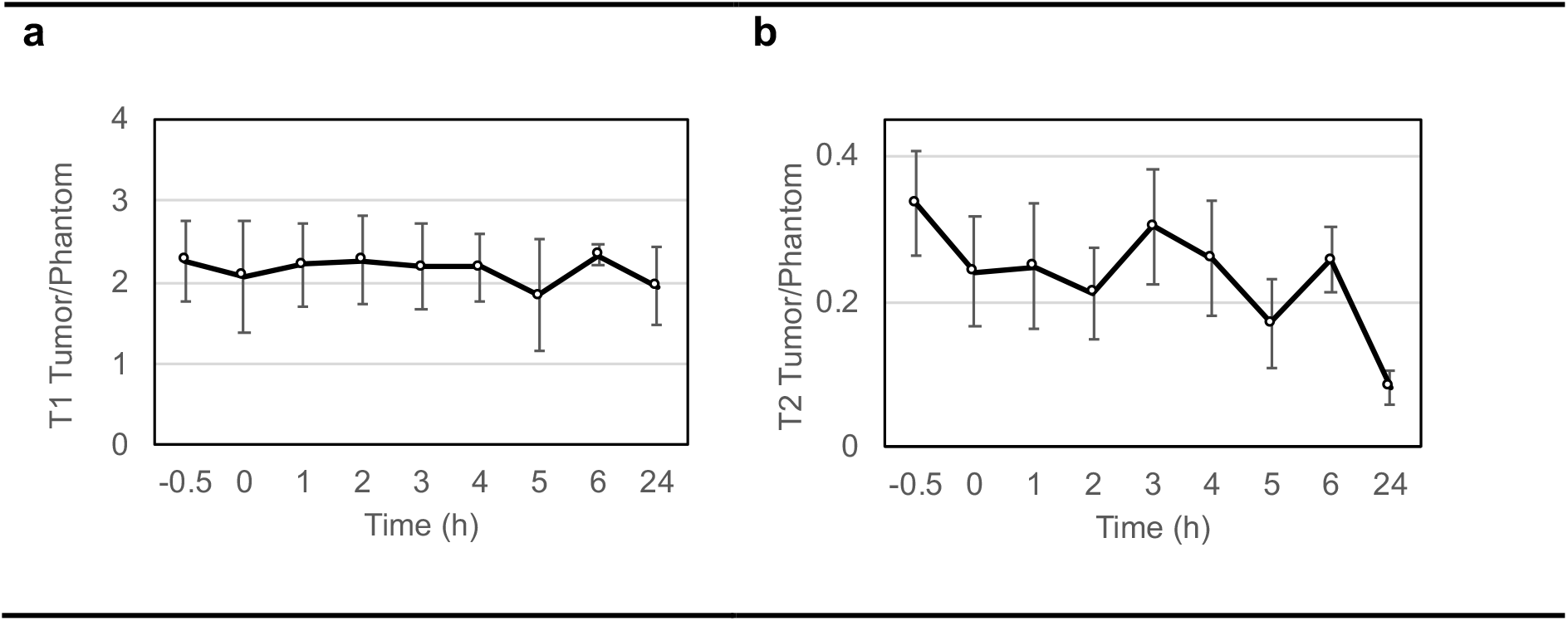
Time-course of *T_1_* and *T_2_* values in xenograft chondrosarcoma tumors before and after intravenous nanoparticle injection. **(a)** Average *T_1_* values in the tumor, normalized to phantom. **(b)** Average *T_2_* values in the tumor, normalized to phantom.

In order to verify the efficiency of our boron-delivery system *in vitro*, CH-2879 chondrosarcoma cells [20] and an ALDH+ radioresistant CSC subpopulation (around 1%) [37] (Figure 5a) were exposed to an epithermal neutron beam at the iBNCT facility [23] (Table S1). Although apoptosis induction after 24h was limited (Figure 5c), BNCT beam exposure resulted in significant DNA damage levels and lower clonogenic survival (Figure 5be). BNCT exposure of cells in the absence of B-MSNs did not trigger DNA damage, suggesting that the biological effects of the neutron beam resulted from BNCT reaction. Comparison of survival curves may allow for a rough estimation of RBEs for dosimetry in this experimental system. Doses resulting in 10% survival (D_10_) were 5.86 Gy for X-rays and 0.42 mA.h (about 6.7×10^11^ n/cm^2^) for neutron beam.

**Figure 5.**
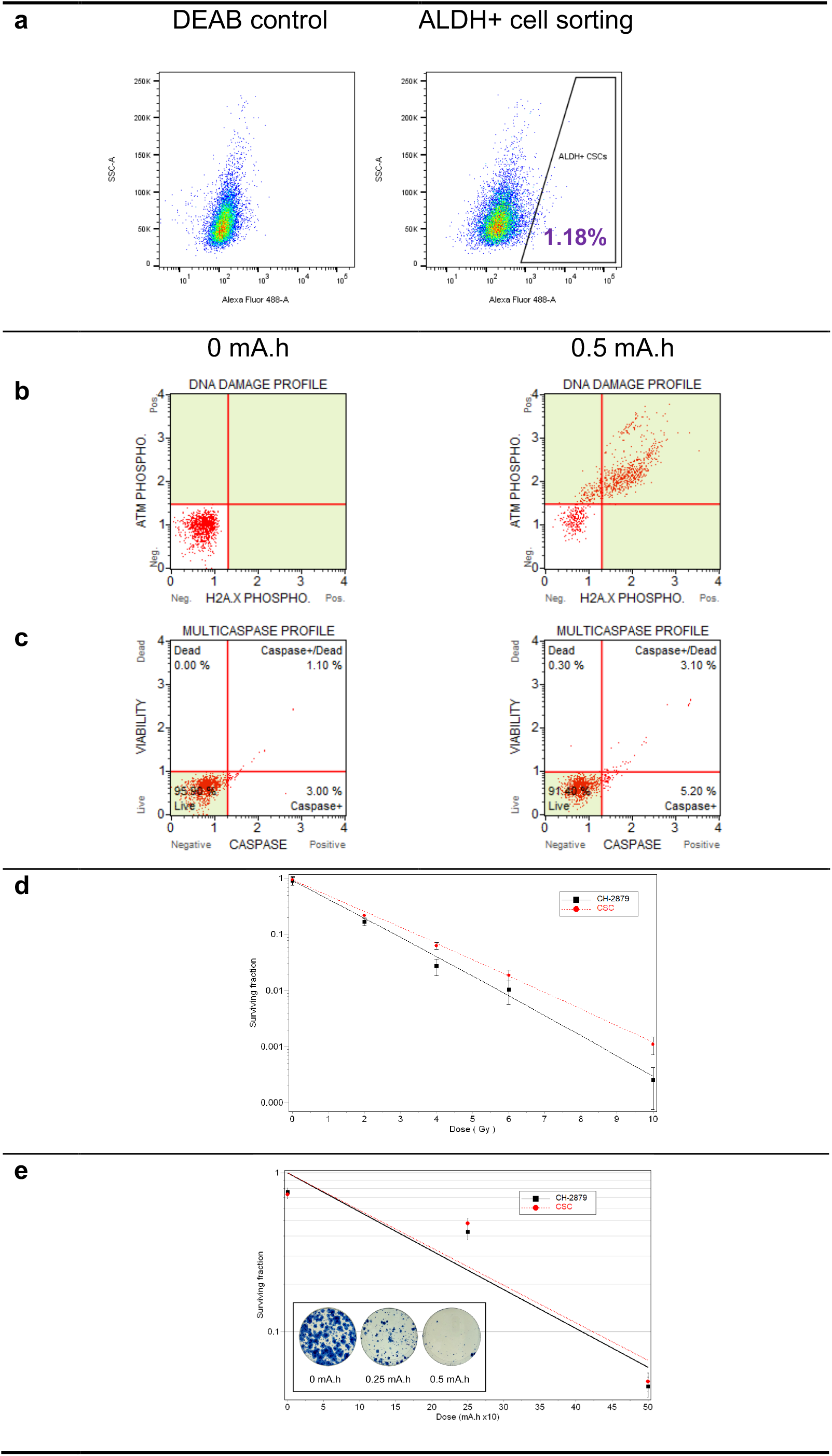
Boron-neutron capture therapy (BNCT) of chondrosarcoma cells with boron-delivery nanoparticles. **(a)** Sorting of chondrosarcoma ALDH+ cancer stem cells. DEAB-treated cells served as negative control. **(b)** BNCT induces DNA damage. ATM and H2AX phosphorylation levels were measured in non-irradiated and irradiated CH-2879 cells, respectively. **(c)** BNCT induces limited apoptosis 24h after irradiation. (d-e) CSCs are comparatively resistant to X-rays **(d)**, but not to BNCT **(e)**, compared with non-CSC CH-2879 cells, as observed after performing clonogenic assays. Neutron beam exposures were performed on cells after administration of mesoporous silica nanoparticles (MSNs) functionalized with activatable cell-penetrating peptide (ACPP).

Interestingly, while CSCs were more resistant to conventional X-ray therapy than the general CH-2879 cell population (Figure 5d), no significant difference was observed in cells exposed to neutron beam (Figure 5e), suggesting that high-LET radiation exposures such as BNCT might be more efficient at targeting CSCs than other treatment modalities, as observed for carbon-ion therapy [7].

## Discussion

The short range of alpha particles generated during the ^10^B(n,α,γ)^7^ Li nuclear reaction of BNCT provides the theorical ability to selectively target tumors cells. This could make BNCT an attractive treatment modality if improvements in boron-delivery and monitoring could be achieved [12]. Only two FDA-approved boron delivery compounds are currently adopted for clinical use: BPA and BSH [38,39]. However, those two compounds lack specificity and do not allow the near-simultaneous measurement of boron biodistribution offered by theranostic delivery systems.

Our multi-functional B-MSNs may be a suitable delivery system for BNCT in some resistant cancers. Other boron-delivery strategies have included the encapsulation of boron-curcumin in poly(lactic-co-glycolic acid) (PLGA) nanoparticles [40], the use of boron cluster-containing polyion complex (PIC) nanoparticles [41], a boron-rich MAC-TAC liposomal system [42], the delivery of dipeptides of BPA and Tyrosine by cancer-upregulated peptide transporters 1 and 2 (PEPT1, PEPT2) [43], or the use of radiolabelled fluoroboronotyrosine (FBY) for Positron Emission Tomography (PET)-guided BNCT [44]. Our B-MSNs exhibit several advantageous features when compared with other organic and inorganic drug delivery systems: well-described synthesis steps and applications, easily tuneable particle and pore size, high flexibility for further functionalization, suitability for theranostic approaches [45]. Furthermore, B-MSNs design and properties may be tailored based on tumor characteristics [46].

Ultimately, clinical dose estimation for BNCT will require both the determination of neutron beam dose components and of boron uptake and biodistribution. Tsukuba-plan, a treatment planning system (TPS) for BNCT has been developed, which employs the Particle and Heavy Ion Transport code System (PHITS) as a Monte Carlo transport code [47,48]. Personalized dosimetry might also be possible by performing real-time detection (using Single Photon Emission Computed Tomography - SPECT) of the gamma-rays emitted by excited ^7^Li as a result of BNCT reactions [49]. The visualization of boron tumor uptake requires the functionalization of boron-delivery systems with a suitable molecular imaging reporter, like ^18^F for PET or Fe/Gd for MRI, while optical imaging may be suitable only for small animal studies. Although PET is usually more sensitive, it requires the use of radioactive tracers, whereas MRI provides high spatial resolution; both techniques are considered sufficiently sensitive for clinical BNCT applications [50]. For example, hepatocytes could be visualized by MRI when cells integrated at least 4 x 10^7^ Gd complexes per cell [51].

Overcoming treatment resistance might require an effective targeting of radioresistant CSCs [52,53]. Therapeutic strategies against CSCs have included inhibition of WNT and NOTCH pathways, ablation using antibody-drug conjugates (ADCs) or epigenetic therapy, each with potential drawbacks or limitations [54]. High-LET radiation treatment, in combination with other targeted therapies, has shown favorable results in bypassing tumor and CSC radioresistance. Using our boron-delivery system, BNCT might be capable of efficiently targeting radioresistant CSCs in hard-to-treat tumors, such as chondrosarcoma. The ability of nanoparticle-based systems to target specific or diffuse tumor sites, as observed in malignant mesothelioma [55], is also of particular interest for BNCT. The combination of different boron-delivery approaches may also improve tumor boron targeting and treatment efficacy [56]. Recently, a number of proton accelerator-based neutron sources have been commissioned for research and clinical use [57], opening new perspectives for the potential development of BNCT as a viable new cancer therapy modality.

## Acknowledgements

This study was funded by OIST and by the R&D Cluster Research Program (Okinawa Prefecture / OIST).

## Abbreviations

ACPP: Activatable cell penetrating peptide
ADC: antibody-drug conjugate
B-MSN: Boron-delivery mesoporous silica nanoparticle
BNCT: Boron neutron capture therapy
BPA: Boronophenylalanine
BSH: Mercaptoundecahydrododecaborate
CSC: Cancer stem cell
CSS: Clear cell sarcoma
DLS: Dynamic light scattering
EPR: Enhanced permeability effect
FITC: Fluorescein isothiocyanate
ICP-MS: Inductively coupled plasma mass spectrometry
LET: Linear energy transfer
MMP: Matrix metalloproteinase
MRI: Magnetic resonance imaging
NP: Nanoparticle
OER: oxygen enhancement ratio
PEG: Polyethylene glycol
PIC: Polyion complex
PLGA: poly(lactic-co-glycolic acid)
RBE: Relative biological effectiveness
TEM: Transmission electronic microscopy

## Notes

#### Summary of Updates

The structure of the manuscript was revised to separate Results and Discussion sections. There is no change in experimental results.

