## Supporting Information for "Functionalized mesoporous silica nanoparticles for innovative boron-neutron capture therapy of resistant cancers"

‡These authors contributed equally

[1] Present Address: Biotage Japan, Koto, Tokyo 136-0071

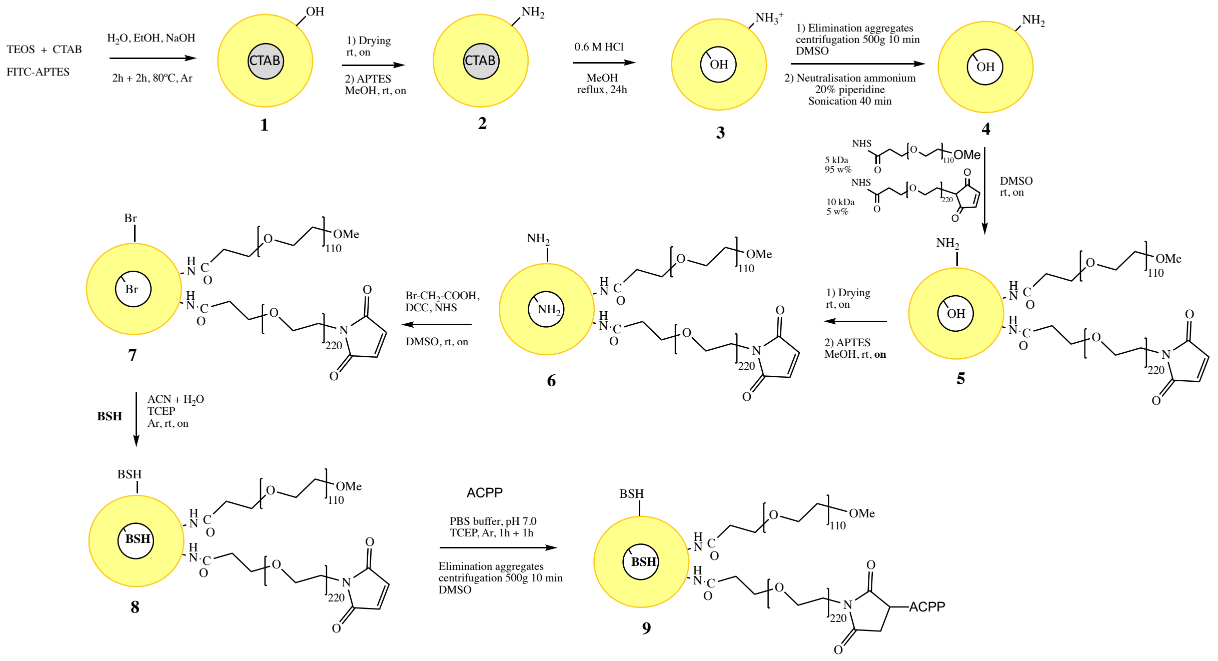

**Scheme S1. Synthesis of multifunctional mesoporous silica nanoparticles.** Synthesis of boron-delivery nanoparticles (B-MSNs).

| Run No. | Beam fluence (mA) | Variance (%) | Dose (mA.h) | Neutron fluence (n/cm$^2$) | Neutron flux (n/cm$^2$/s) |
|---|---|---|---|---|---|
| 1 | 1.28 | 6.62 | 0.25 | $3.8x10^{11}$ | $5.3x10^{8}$ |
| 2 | 1.28 | 3.75 | 0.5 | $7.6x10^{11}$ | $5.3x10^{8}$ |
| 3 | 1.28 | 4.97 | 0.5 | $7.6x10^{11}$ | $5.3x10^{8}$ |
| 4 | 1.29 | 6.14 | 0.5 | $7.7x10^{11}$ | $5.3x10^{8}$ |
| 5 | 1.29 | 6.72 | 0.25 | $3.9x10^{11}$ | $5.3x10^{8}$ |
| 6 | 1.28 | 8.62 | 0.5 | $7.6x10^{11}$ | $5.3x10^{8}$ |
| 7 | 1.3 | 4.87 | 0.5 | $7.8x10^{11}$ | $5.3x10^{8}$ |
| 8 | 1.3 | 4.27 | 0.5 | $7.7x10^{11}$ | $5.3x10^{8}$ |

**Table S1. Neutron beam characteristics.**

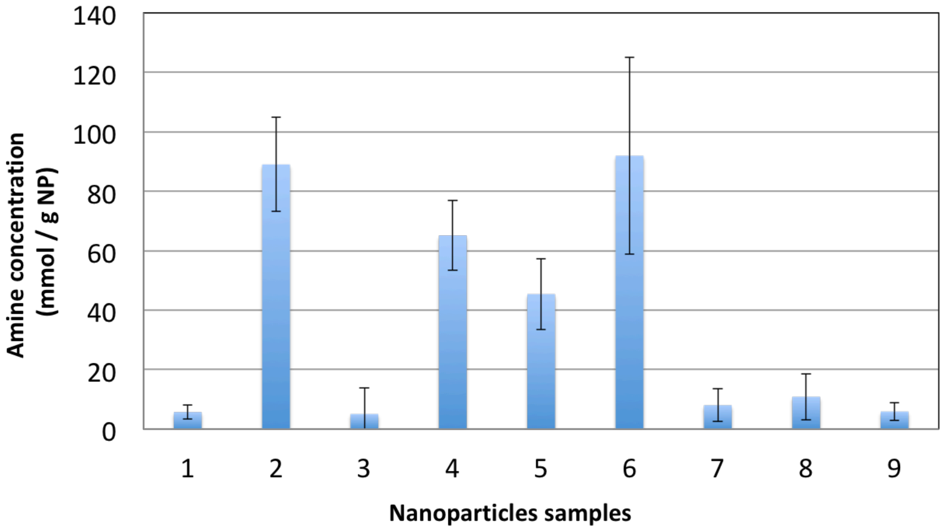

**Figure S1. Amine quantification.** Amine concentration was measured at each B-MSN synthesis step by using the Fmoc method.

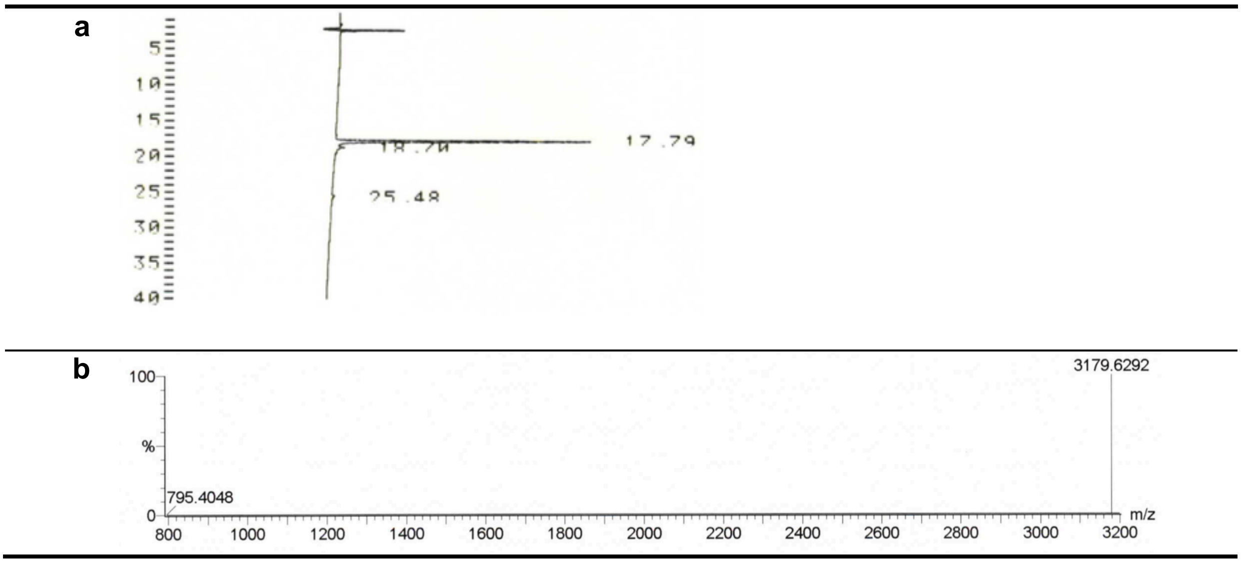

**Figure S2. Purification and characterization of activatable cell penetrating peptide (ACPP).** (a) Reversed-phase HPLC chromatogram. (b) High-resolution mass spectrometry (HR-MS) spectrum.

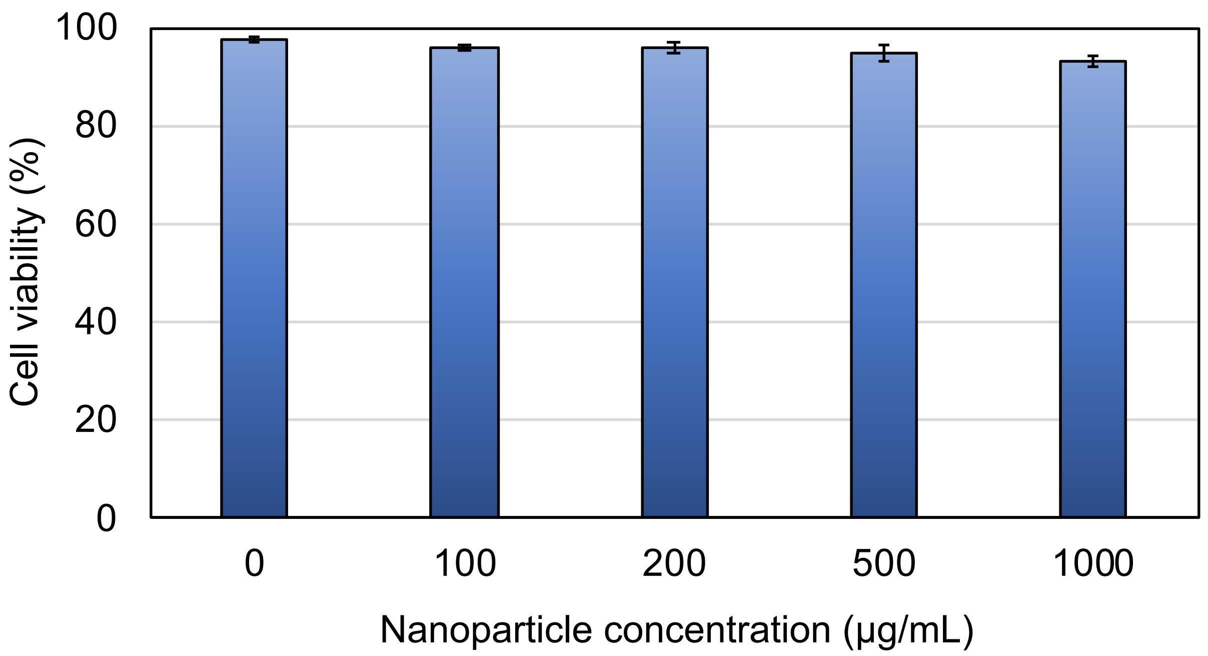

**Figure S3**. **Verification of *in vitro* nanoparticle toxicity.** Cell viability was measured 48h after B-MSN administration at increasing concentrations.

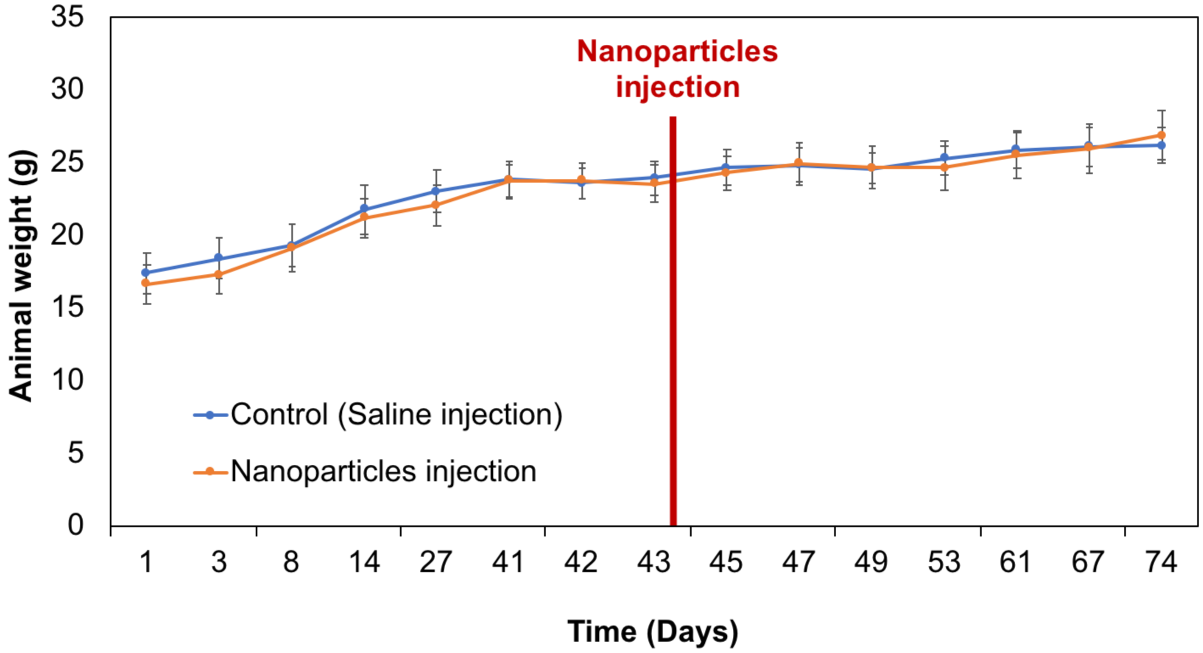

**Figure S4. Verification of *in vivo* nanoparticle toxicity.** Animal weight was measured for a month after tail-vein administration of nanoparticles.

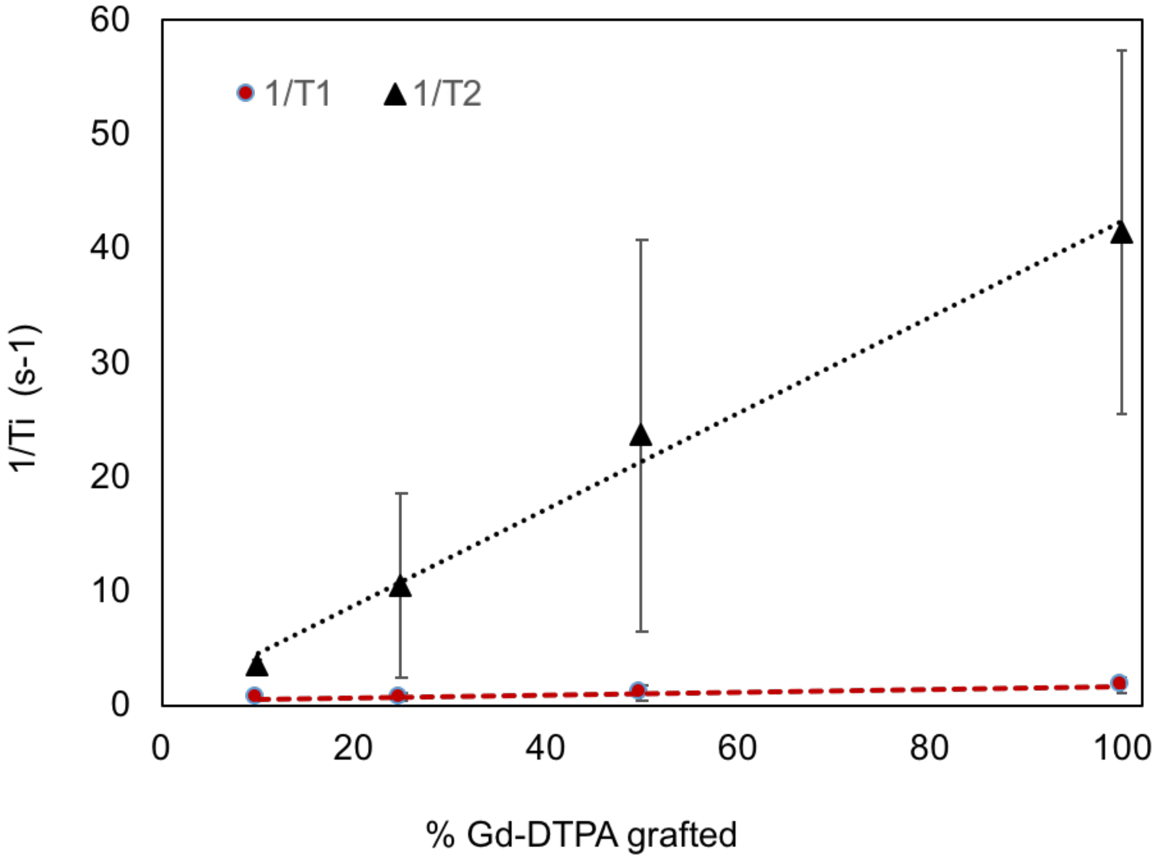

**Figure S5. T1 and T2 values in Magnetic Resonance Imaging (MRI) as a function of MSN Gadolinium loading on nanoparticles.**

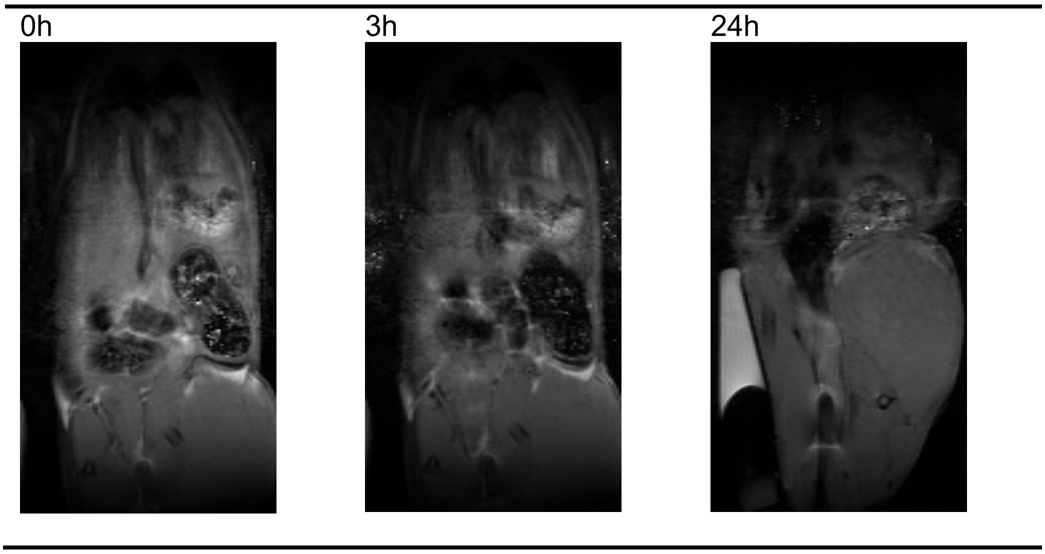

**Figure S6. T2-weighted MRI images of tumor-bearing nude mice after tail vein administration of gadolinium-nanoparticles.**

### Supplementary Material and Methods

**Amine quantification.** Aminated nanoparticles (20 mg) were dispersed in 5mL dimethylformamide (DMF), then 100 mg 9-Fluorenylmethoxycarbonyl chloride (Fmoc-Cl) was added. The mixture was stirred overnight at room temperature, centrifugated, washed with methanol and finally dried overnight. In a second step, Fmoc-protected NPs (5.0 mg) were suspended in a mixture of DMF/piperidine (5.0 mL, 4/1) and bath sonicated for 40 min. The suspension was centrifugated and the absorbance of the solution was measured by UV-vis spectrophotometry at $\lambda$ = 300 nm ($\varepsilon$ = 7800 $mol^{-1}.L.cm^{-1}$) [1].

**PEG quantification**. PEGylated B-MSNs (5.0 mg) were suspended in a solution made of Sodium deuteroxide (NaOD) (200 mM) and DMF as internal standard (1 mM) in Deuterium oxide (D2O) ($V_{tot}$ = 500 µL). The dissolution reaction was stirred over night at room temperature and analyzed by Proton nuclear magnetic resonance ($^1$H NMR) using a 400 MHz JNM-ECZ400S/L1 NMR (JEOL, Tokyo Japan) through the integration of the HCOOH (1H) ($\delta$ = 8.3 ppm) and $CH_2$-$CH_2$-O- (4H) ($\delta$ = 3.7 ppm) peaks.

**Evaluation of nanoparticles cytotoxicity.** *In vitro* viability was measured by trypan blue exclusion, using a TC10 automated cell counter (Bio-Rad). To evaluate *in vivo* toxicity, nanoparticles were injected in the tail vein of healthy mice and body weight was measured for several weeks.
